# DTI-Voodoo: machine learning over interaction networks and ontology-based background knowledge predicts drug–target interactions

**DOI:** 10.1101/2021.04.28.441733

**Authors:** Tilman Hinnerichs, Robert Hoehndorf

## Abstract

**Motivation:** *In silico* drug–target interaction (DTI) prediction is important for drug discovery and drug repurposing. Approaches to predict DTIs can proceed indirectly, top-down, using phenotypic effects of drugs to identify potential drug targets, or they can be direct, bottom-up and use molecular information to directly predict binding potentials. Both approaches can be combined with information about interaction networks.

**Results:** We developed DTI-Voodoo as a computational method that combines molecular features and ontology-encoded phenotypic effects of drugs with protein–protein interaction networks, and uses a graph convolutional neural network to predict DTIs. We demonstrate that drug effect features can exploit information in the interaction network whereas molecular features do not. DTI-Voodoo is designed to predict candidate drugs for a given protein; we use this formulation to show that common DTI datasets contain intrinsic biases with major affects on performance evaluation and comparison of DTI prediction methods. Using a modified evaluation scheme, we demonstrate that DTI-Voodoo improves significantly over state of the art DTI prediction methods.

**Availability:** DTI-Voodoo source code and data necessary to reproduce results are freely available at https://github.com/THinnerichs/DTI-VOODOO.

**Supplementary information:** Supplementary data are available at https://github.com/THinnerichs/DTI-VOODOO.

## 2 Introduction

Identifying drug–target interactions (DTIs) is a crucial step in drug discovery; finding novel DTIs for approved drugs can be used for drug repurposing, either by finding new drugs for a known target or finding a drug for a novel target involved in a disease process. Inferring the interactions between drugs and their targets can help to analyze and identify potential desired or adverse drug effects as well as desirable therapeutic effects. While *in vitro* DTI prediction is time consuming, computational *in silico* DTI predictors can screen for millions of interactions within a short time. Determining DTIs computationally can therefore help to mitigate the costs and risks of drug development.

Computational methods are widely applied to predict DTIs and many computational methods have been developed. These methods can be broadly classified into “top-down” and “bottom-up” approaches. Top-down approaches start from observable characteristics resulting from a drug–target interaction, such as side-effects or the diseases treated by a drug, and infer likely molecular mechanisms (i.e., the interaction) using these observations. Bottom-up approaches start from molecular features such as molecular structure or fingerprints associated with drug and protein, and predict interactions from this information.

Both bottom-up and top-down approaches to drug–target interaction prediction bear some advantages and limitations. Generally, bottom-up methods face the challenge to predict whether a chemical structure binds to a protein given their molecular properties; whether two entities interact depends not only on the molecular structure of the entities (where binding sites and molecular forces need to be determined for accurate prediction) but also properties such as in which celltypes and anatomical structures a protein is expressed. Top-down methods use information about physiological effects of drugs for DTI prediction, such as side-effect similarity (Campillos *et al*., 2008), that is largely complementary to knowledge gained from molecular properties. While methods that rely on molecular information are directly predicting whether two molecules could interact, top-down methods base on more indirect means and infer a DTI from observable effects resulting from the interaction.

Both approaches may be used in conjunction with network inference (Chen *et al*., 2015). Biological networks used for DTI prediction include protein–protein interaction networks (Feng *et al*., 2017; Lee and Nam, 2018) and networks including several other types of biological relations, including similarity between represented entities (Ding *et al*., 2013; Gottlieb *et al*., 2011). Network-based DTI prediction methods use the guilt-by-association principle (Oliver, 2000) and assume that a protein is a likely target for a drug if many of the protein’s neighbors in the interaction network are targets of the drug (Gillis and Pavlidis, 2012). Network-based methods have been applied successfully to DTI prediction. However, if DTIs are taken as direct physical interactions between a drug and protein, it remains an unresolved question whether the network-based guilt-by-association hypothesis is true, or whether an interaction of a drug and protein dysregulates several of the protein’s interaction partners, and therefore resulting in effects that are not direct interactions but only downstream consequences of an interaction.

Progress in machine learning using graph neural networks can allow us to test this hypothesis and combine both bottom-up and top-down features with a network in a single machine learning model. In particular, Graph Convolutional Networks (Kipf and Welling, 2016) and their variants operate on different types of kernels (Defferrard *et al*., 2016; Bianchi *et al*., 2019), including attention mechanisms (Veličković *et al*., 2017), and different forms of exploring node neighborhoods (Klicpera *et al*., 2018; Hamilton *et al*., 2017) can combine different types of features and graph-based information. They have previously been applied for a number of tasks, including prediction of protein functions (Zitnik and Leskovec, 2017), cancer drug response (Liu *et al*., 2020) and drug–target affinity prediction (Nguyen *et al*., 2020).

Potential biases resulting from the underlying datasets (Pahikkala *et al*., 2014) which may affect model evaluation and comparison pose a challenge for DTI prediction. Firstly, novel drugs are often developed by altering non-functional components of a drug, leading to two and more very similar drugs designed to target the same proteins (Overington *et al*., 2006). This can result in a bias when it leads to hidden duplicates or highly similar compounds that are distributed among training and evaluation dataset, resulting in a better (measured) predictive performance than it would be expected when the model is applied to identify drugs that target a protein for which no drugs yet exist. Secondly, some proteins (which we call *hub proteins*) have significantly more known interactions with drugs than others. In the STITCH database, 5% of the proteins have 40% of the interactions, and similar distributions are present in other datasets (Wishart *et al*., 2007, 2017); preferentially predicting these proteins may increase predictive performance while not reflecting the actual performance when applied to a new protein (i.e., a protein for which no interactions are known). These differences in the number of drugs targeting certain proteins may be the result of study bias where more “valuable” proteins have more drugs designed to target them due to their involvement in more common diseases (or diseases for which drugs can be more profitably marketed). This might affect common evaluation schemes where it is possible to exploit these biases within DTI prediction (Wang and Kurgan, 2018). van Laarhoven and Marchiori (2014) showed that several bias can be exploited on the dataset of Yamanishi *et al*. (2008).

We developed DTI-Voodoo as a method for predicting DTIs. We use an ontology-based machine learning method (Chen *et al*., 2020) to encode phenotypic consequences of DTIs and deep learning methods to encode molecular features. We combine both using a protein interaction network which we exploit with the aid of a graph neural network. We use this model to test whether molecular or phenotype features benefit from the network information and find that only phenotype features localize on the graph whereas molecular features do not. We further evaluate and compare DTI-Voodoo against several DTI prediction methods and demonstrate a substantial improvement of DTI-Voodoo over the state of the art in predicting drugs that target a protein. We also identify and characterize several biases in both training and evaluating DTI prediction methods, and make recommendations on how to avoid them. DTI-Voodoo is available as Free Software at https://github.com/THinnerichs/DTI-VOODOO.

## 3 Methods

### 3.1 Problem Description

DTI-Voodoo aims to solve the following problem: for a given drug and a given protein we want to determine whether those interact or not. We do not differentiate between types of interaction such as activation and inhibition, and do not predict the strength of the interaction. We treat all drug–protein pairs without a known interaction as negatives and therefore formulate the problem as a binary classification task.

### 3.2 Datasets

We obtain a dataset consisting of 12,884 human proteins with 340,627 links from STRING (Szklarczyk *et al*., 2014). For the drug-target interactions, we use 229,870 links from the STITCH database (Szklarczyk *et al*., 2015). As both STRING and STITCH provide confidence scores for each association, we filtered them as advised by a threshold of 700, therefore retaining only high-confidence interactions.

We utilize the PhenomeNET ontology (Hoehndorf *et al*., 2011), an ontology integrating ontologies such as the Human Phenotype Ontology (Köhler *et al*., 2018), Gene Ontology (Ashburner *et al*., 2000; Carbon *et al*., 2020), Mammalian Phenotype Ontology (Smith and Eppig, 2009) and several others. We obtained side effects and their links to drugs from SIDER (Kuhn *et al*., 2015); SIDER contains side effects encoded using identifiers from the MedDRA database (Mozzicato, 2009). We mapped side effects to the PhenomeNET ontology using the *Phenomebrowser*.*net*, which provides a SPARQL query endpoint for the mentioned resources. The overall structure is shown in Supplementary Figure 1.

For comparative evaluation, we use the gold standard dataset introduced by Yamanishi *et al*. (2008) consisting of 1,923 interactions between 708 drugs and 1,512 proteins, and the BioSnap dataset (Zitnik *et al*., 2018) which consists of 5,017 drug nodes, 2,324 gene nodes and 15,138 edges.

We only use proteins in our analysis that have at least one link in STRING or one association in PhenomeNET, and drugs with at least one side effect. Therefore, the intersection between these resources yields 1,428 drugs and 7,368 human proteins with 32,212 interactions for STITCH, 1,837 interactions between 680 drugs and 1,458 proteins for Yamanishi, and 6,498 links between 949 drugs and 2,221 proteins for BioSnap dataset. We provide links to and methods for obtaining and processing the necessary data in our Github repository.

### 3.3 Model

Our model combines “top-down” and “bottom-up” information for drug– target identification. We consider an approach to be “top-down” when observable characteristics of either a drug (such as a drug effect) or protein (such as a protein function, or phenotypes resulting from a loss of function) are used to provide information about a molecular mechanisms; we consider an approach “bottom-up” when structural or other molecular information is used to determine a mechanism. In order to build a method that incorporates both top-down and bottom-up features, we first create a model for each type of feature separately. As features for the bottom-up model, we use features derived from molecular structures of drugs from the *SmilesTransformer* (Honda *et al*., 2019) and molecular features for proteins from *DeepGOPlus* (Kulmanov and Hoehndorf, 2019). *SmilesTransformer* is an autoencoder trained over the SMILES strings, and therefore captures (some aspects of) the molecular organization of each drug in an unsupervised manner. *DeepGOPlus* provides features derived from protein amino acid sequences which are useful to predict protein function.

As phenotypes and functions are encoded through ontologies, we use DL2Vec (Chen *et al*., 2020) to obtain ontology based representations for use as top-down features. DL2vec constructs a graph by introducing nodes for each ontology class and edges for ontology axioms, followed by random walks starting from each node in the graph. These walks are encoded using a Word2vec (Mikolov *et al*., 2013) model. Therefore, DL2Vec generates representations that enable to encode drug effects or protein functions while preserving their semantic neighborhood within that graph.

#### 3.3.1 Half-twin neural networks and feature transformation

As we want to learn from the similarity of drug side effects and protein phenotypes, we use a deep half-twin neural network with a contrastive loss using cosine similarity. A half-twin neural network aims to learn a similarity between two embeddings of variable but same dimension. As the original feature space may have varying dimensionality, we first process them using a fully connected neural network layer which takes as input an embedding and outputs a representation of a particular size, i.e., we use this layer as a trainable feature transformation and apply it to reduce the representation size of the embeddings for drugs and proteins separately. An example structure for both types of features can be found in Supplementary Figure 2. The use of this trainable feature transformation layer enables flexible experimentation as both ontology and molecular feature for both drugs and proteins are reduced to the same dimensionality for varying sizes of inputs; this allows for a high amount of modularity across different experimental setups by adding different kinds of features into the model. Additionally, the generated features may be used for other tasks. We follow the results of *DL2vec* (Chen *et al*., 2020) and use *σ* := LeakyReLU as activation function which leads to improved performance compared to other activation functions.

#### 3.3.2 Graph convolutional layers

We include these molecular and ontology-based sub-models within a graph neural network (GNN) (Kipf and Welling, 2016). The graph underlying the GNN is based on the protein–protein interaction (PPI) graph. The PPI dataset is represented by a graph *G* = (*V, E*), where each protein is represented by a vertex *v ∈ V*, and each edge *e ∈ E* ⊆ *V × V* represents an interaction between two proteins. Additionally, we introduce a mapping *x* : *V* → ℝ^*d*^ projecting each vertex *v* to its node feature *x*_*v*_ := *x*(*v*), where *d* denotes the dimensionality of the node features.

A graph convolutional layer (Kipf and Welling, 2016) consists of a learnable weight matrix followed by an aggregation step, formalized by

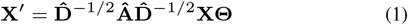

where for a given graph *G* = (*V, E*), Â= *A* + *I* denotes the adjacency matrix with added self-loops for each vertex, *D* is described by 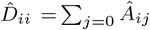, a diagonal matrix displaying the degree of each node, and Θ denotes the learnable weight matrix. Added self-loops enforce that each node representation is directly dependent on its own preceding one. The number of graph convolutional layers stacked equals the radius of relevant nodes for each vertex within the graph.

The update rule for each node is given by a message passing scheme formalized by

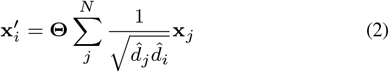

where both 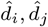 are dependent on the edge weights *e*_*ij*_ of the graph. With simple, single-valued edge weights such as *e*_*ij*_ = 1 *∀*(*i, j*) *∈ E*, all 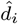 reduce to *d*_*i*_, i.e., the degree of each vertex *i*. We denote this type of graph convolutional neural layers with GCNConv.

While in this initial formulation of a GCNConv the node-wise update step is defined by the sum over all neighboring node representations, we can alter this formulation to other message passing schemes. We can rearrange the order of activation function *σ*, aggregation AGG, and linear neural layer MLP with this formulation as proposed by Li *et al*. (2020):

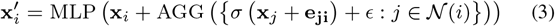

where we only consider *σ ∈* {ReLU, LeakyReLU}. We denote this generalized layer type as GENConv following the notation of PyTorch Geometric (Fey and Lenssen, 2019). While the reordering is mainly important for numerical stability, this alteration also addresses the vanishing gradient problem for deeper convolutional networks (Li *et al*., 2020). Additionally, we can also generalize the aggregation function to allow different weighting functions such as learnable SoftMax or Power for the incoming signals for each vertex, substituting the averaging step in GCNConv. Hence, while GCNConv suffers from both vanishing gradients and signal fading for large scale and highly connected graphs, each propagation step in GENConv emphasizes signals with values close to 0 and 1. The same convolutional filter and weight matrix are applied to and learned for all nodes simultaneously. We further employ another mechanism to avoid redundancy and fading signals in stacked graph convolutional networks, using residual connections and a normalization scheme (Li *et al*., 2019, 2020) as shown in Supplementary Figure 3. The residual blocks are reusable and can be stacked multiple times.

#### 3.3.3 Combined prediction model

Combining half-twin and graph convolutional neural networks, we map all protein representations to their respective node features, initializing the graph convolutional update steps. The resulting representations are used for a similarity prediction. When combining ontology and molecular features with or without the graph model, we concatenate both protein features and both drugs features, before plugging them into the graph model for the similarity computation. An overview of the model architecture, combining both feature types, is shown in Supplementary Figure 4. Here the original representations are transformed by a dense layer and then used as input of a stack (with height 3) of residual graph convolutional blocks.

#### 3.3.4 Hyperparameter tuning

As the number of drug-targets are sparse with respect to the amount of both drugs and proteins considered, the training, validation and testing datasets are imbalanced. As there are only 22, 336 links in the considered STITCH subset, the ratio

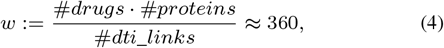

consequently needs suitable compensation in the computed loss function and appropriate metrics for the evaluation.

Therefore, we weight all positive drug-protein pair samples with this ratio by introducing the following loss function with respect to binary cross-entropy:

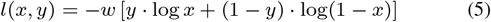

for a given prediction *x* and target *y*, and positive weight *w* defined by equation (4). We average this loss among all drug-protein pairs in the training set, leading to a stable environment for the *Adam* optimization algorithm (Kingma and Ba, 2015). We implemented a 5-fold cross validation split among the proteins. Furthermore, we used early stopping in the training process.

To find the best hyperparameter configuration for the proposed model, we performed a grid search to find the most expressive and non-redundant representation. We pretrained the bottom-up and the top-down model separately and aimed at best performing models with respect to our evaluation metrics. We optimized embedding sizes, depth of the neural network, optimizer, learning rate and layer types using an extensive, manual grid search. Starting from shallow feature transformations with an embedding size of 10, we scaled the network up to residual structures with up to 10 hidden layers leading to embeddings of size 4000, testing different network widths and learning rates for each configuration.

### 3.4 Evaluation and metrics

#### 3.4.1 Splitting schemes

As DTI prediction is dependent on both drugs and proteins, there are multiple ways of determining training, validation and testing sets of pairs to evaluate each model. For cross-validation, we can perform the split over DTI pairs, over drugs and over proteins, respectively; when splitting over drugs or proteins, the entities (drugs or proteins) are separated and all their associations included in the split. Only when splitting by protein or drug are unseen entities guaranteed to be shown to the model in the validation and testing phase. The models resulting from the different splitting schemes may have different expressiveness and exploit different information in DTI prediction, as different information is known in the training and testing phase.

#### 3.4.2 Metrics

To assess each model, we compute a variety of common metrics for binary classification. As the datasets are highly imbalanced, we use the area under the receiver operating characteristic curve (AUROC) on training, validation and testing split.

We calculate the AUROC by computing true positive rate at various false positive rate thresholds and use trapezoidal approximations to estimate the area under the curve. We refer to this measure as MacroAUC.

We also calculate the MicroAUC score. For given lists *D* and *P* of drugs and proteins, respectively, and a set of known interactions *Int* := {(*d*_*i*_, *p*_*i*_)}, *MicroAUC* is calculated as the average per entity *AUROC* score. For example, the protein-centric score can be formalized as: given labels *l* : *D × P* → {0, 1} and predictions *y* : *D × P* → [0, 1], we define

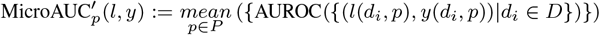

In some cases, the *MicroAUC* score may not be defined as in some datasets some proteins or drugs have no interactions, leading to an infeasible *T PR* = 0 for all thresholds and an undefined *AUROC* score for that entity. As this is quite common for DTI datasets, we do not omit but impute the *MicroAUC* interpolating linearly for those entities using the accuracy for this subset:

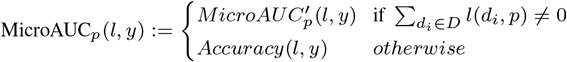

Drugs and proteins can be interchanged in this formulation. We refer to the different measures as protein-centric microAUC (*MicroAUC*_*p*_) and a drug-centric microAUC (*MicroAUC*_*d*_). We further compare MicroAUC_*p*_ with imputation and 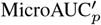 without imputing undefined *MicroAUC* values (but omitting them) in Supplementary Table 1.

Further, we choose the AUROC over other measures such as the area under precision recall curve (AUPRC) as primary metric to compare different methods; AUPRC is sensitive to data imbalances (Jeni *et al*., 2013) and therefore more challenging to apply to comparing different DTI prediction methods.

## 4 Results

### 4.1 DTI-Voodoo: computational model to identify drugs that target a protein

We developed DTI-Voodoo as a computational model to predict drug– target interactions. Specifically, given a protein, DTI-Voodoo will identify and rank drugs that likely target this protein. DTI-Voodoo combines two types of features: structural information for drugs and proteins that can be used to determine if the drug and protein physically interact, and information about phenotypic effects of drugs and changes in protein function that may “localize” on an interaction network (i.e., neighboring nodes will share some of these features or are phenotypically similar). As structural features, DTI-Voodoo uses structural representations of drugs from the SMILES transformer (Honda *et al*., 2019) and representations of protein amino acid sequences from DeepGOPlus (Kulmanov and Hoehndorf, 2019). DTI-Voodoo learns representations of drug effects and protein functions using the ontology-based machine learning method DL2Vec (Chen *et al*., 2020) and ontology-based annotations of drugs and proteins.

We construct a graph with proteins as nodes and protein-protein interactions as edges, mapping the protein features to each target as node features. DTI-Voodoo then propagates information among the PPI network utilizing graph convolutional steps, calculates the similarity of drug and protein representations, and predicts whether there is an interaction. The full workflow scheme is depicted in Figure 1

**Figure 1.**
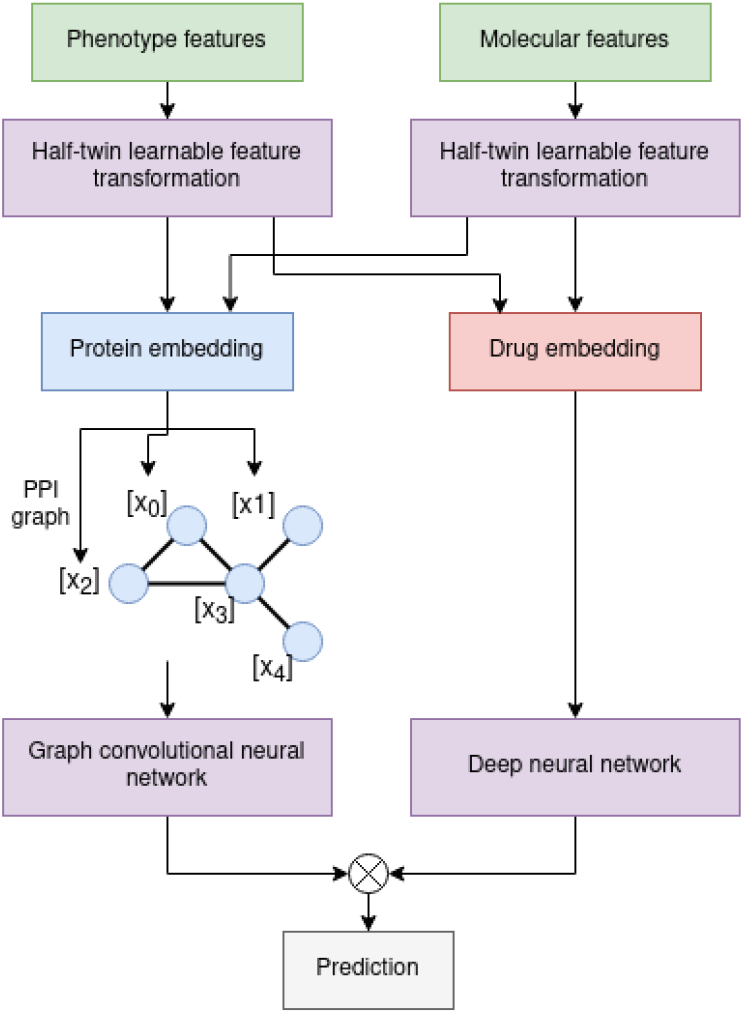
Full DTI prediction model based on the pretrained learnable feature transformations (LFT) for either molecular structure or ontology based features. The transformed protein representations are added to each corresponded protein as node features for the graph convolutional steps.

We evaluate our model’s ability to identify drug–target interactions using different approaches and datasets. First, we perform a cross-validation over proteins and validate our results. A cross-validation over proteins aims to evaluate how the model performs when tasked to identify drugs that may target a “novel” protein, i.e., one not seen during training, or a protein for which a drug that targets it should be predicted.

We trained, validated and finally tested all considered models on the STITCH dataset using a 5-fold cross-validation over a protein split; we then selected the best-performing models (with respect to *MicroAUC*_*p*_, see Section 3.4.2), and retrain them from scratch in a 5-fold protein-split cross-validation on the Yamanishi benchmark dataset to avoid validation overfitting and yield more realistic testing results. To evaluate the influence of the different features separately, and to determine whether they “localize” on the PPI graph (and therefore can be exploited successfully by the graph neural networks), we train and evaluate models with different types of features, and with and without inclusion of the PPI graph, separately. We comparing the molecular (MolPred) and phenotype-based (OntoPred) prediction model, and a combination of both where we concatenate both types of features. Table 1 shows the results of these experiments.

**Table 1.**
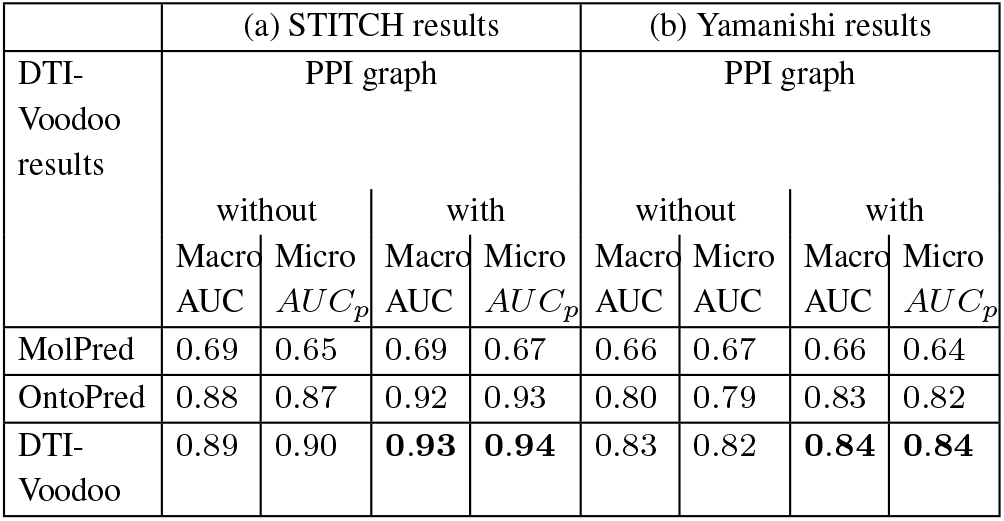
Results for DTI-Voodoo on the STITCH and Yamanishi datasets evaluated with 5-fold cross-validation. We call the model using only molecular features MolPred and the model using only ontology-based features OntoPred. DTI-Voodoo combines both types of features.

We find that the model using ontology-based features (*OntoPred*) is showing better performance on STITCH compared to using only molecular features. We also observe that only the model using ontology-based features results in increased performance when incorporating the PPI graph. This increase can be observed with different graph neural network architectures and configurations (Supplementary Table 2). While the *GCNConv* and *GENConv* architecture already shows some minor improvement, the use of *ResGraphConv* results in larger performance improvements. *ResGraphConv* blocks add a large amount of additional learnable parameters to the network, leading to more expressive power. To test whether the observed improvement is due to the number of learnable parameters added or the result of better exploiting the information about PPIs, we experiment with a graph model in which all graph convolutional neural layers in the residual blocks are removed, resulting in a model with similar parameters but without the ability to use graph-based information. This pruned network, with no information on the protein–protein interactions, reached very similar results to the original *OntoPred* model and showed no improvement (Supplementary Table 2).

The improvements when including the graph are only provided by the *GENConv* graph convolution scheme which includes the *ResGraphConv* blocks; *GCNConv* and other graph convolutional methods fail to achieve any gain in comparison to the plain *OntoPred* performance even when combined with the residual blocks. The discrepancy between *GENConv* and other graph convolutional methods may be the result of numerical instability and fading signals (Li *et al*., 2020).

Our results demonstrate that the inclusion of graph information can increase performance when ontology-based features are used but not when molecular features are used alone. This observation allows us to conclude that information about protein functions localizes on the graph whereas molecular features do not.

We further investigate the performance on the specific interaction types *inhibition* and *activation* that are available within the STITCH database. The results are summarized in Supplementary Table 3; we find no difference between the performance on specific interaction types and the complete dataset where we do not separate types of interaction.

### 4.2 Protein-centric evaluation

The goal of DTI-Voodoo is to find candidate drugs that target a specific protein; however, so far, we do not evaluate this application but rather how DTI-Voodoo would perform in finding plausible drug–target interactions among all possible interactions (since we use the MacroAUC as our main evaluation measure). This evaluation does not correspond to the application of DTI-Voodoo in finding drugs that target a specific protein. To provide a better estimate on how DTI-Voodoo performs for individual protein targets, we use micro-averages between proteins and compute the *MicroAUC* (see Section 3.4.2); to determine *MicroAUC*, we average the performance (true and false positive rates) per protein instead of across all drug–protein pairs; the resulting measure can therefore better estimate how DTI-Voodoo performs when tasked with finding a drug that targets a specific protein.

Furthermore, we hypothesize that it may be possible for a machine learning model to exploit biases in drug–target interaction data to achieve relatively high prediction performance without obtaining a biologically meaningful signal. For example, hub proteins may have a large number of interactions, or certain drugs interact with many proteins, and preferentially predicting these interactions may increase predictive performance even in the absence of any biological features. To test this hypothesis, we design a “naïve” baseline model that predicts the same list of proteins for each drug based only on the number of known drug–target interactions for a protein. Specifically, given lists *D* and *P* of drugs and proteins and a set of known interactions ℐ := {(*d*_*i*_, *p*_*i*_)}, we construct an interaction matrix *M*_*int*_ *∈* {0, 1}^|*D*|*×*|*P* |^ with

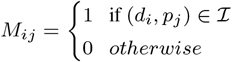

describing for all drug–protein pairs whether there is a known interaction or not. We now rank all proteins *p*_*j*_ *∈ P* descending by their number of drug interactors by summing over the columns of *M*_*ij*_ and ranking these sums:

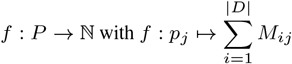

Our “naïve” predictor *P*_*k*_ predicts all drugs to interact with the top *k* targets with respect to the introduced ranking:

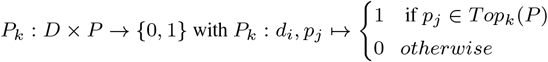

with the only hyperparameter *k*.

The prediction *P*_*k*_ is not dependent on the drug *d*_*i*_ and will predict the same ranked list of drugs for all proteins; consequently, this naïve predictor does not rely on any biological features and will not predict any novel information about interactions between drugs and proteins; the naïve predictor only exploits imbalances in the evaluation set to make predictions that may perform well. The way in which we formulated the naïve predictor, it is not applicable for a protein split cross-validation as the number of interactions for each protein in the validation set is unknown.

We apply this naive predictor on both the STITCH and Yamanishi datasets, using the full datasets as well as a 5-fold cross-validation over drugs and over drug–protein pairs to compare the prediction results directly to DTI-Voodoo. For each fold, we gradually increase *k* to determine the best performance for each fold. Using the full dataset, drug–target split, and drug split, we obtain the following MacroAUC results: for the STITCH database, we obtain a performance of 0.76 on the whole dataset, 0.70 for the drug–target pairs and 0.73 in case of the drug splitting scheme; on the Yamanishi dataset, we obtain MacroAUC scores of 0.88, 0.84 and 0.85, for the total dataset, drug–target pair and drug split, respectively. The naïve predictor shows higher performance on the Yamanishi dataset than on STITCH, and a substantial gain in comparison to an expected random predictor on both datasets. In the following, we utilize this 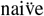 predictor as baseline to compare its performance to state of the art models and DTI-Voodoo.

For comparison with the state of the art methods, we chose the best performing methods for drug–target interaction prediction that were previously evaluated on the Yamanishi benchmark dataset. These methods include DTIGEMS+ (Thafar *et al*., 2020) and DTI-CDF (Chu *et al*., 2019) which have showed superior results in comparison to numerous works. Furthermore, we added DTINet (Luo *et al*., 2017) as method for comparison which has been used to develop a number of methods such as NeoDTI (Wan *et al*., 2019) with similar methodology.

We evaluate all models on their recommended splitting scheme choice, hyperparameters and folds in cross-validation, measuring their respective AUROC. We further evaluate each model by performing a protein-wise cross-validation determining the MacroAUC and *MicroAUC*_*p*_. For this evaluation, we allow sub-sampling of negatives for the training process but not for the validation and testing phase as real world applications of these models would have to deal with possibly imbalanced data.

The results of our experiments are summarized in Table 2; we calculated the performance of all compared methods over their original splitting scheme and over a protein split. We find that there is a large difference in performance when evaluating over a drug–target pair split compared to a protein split, with generally higher performance achieved when using the drug–target pair split. Second, when evaluating the same methods over a protein split, we find a substantial performance difference in comparison to the splitting scheme used in the original evaluation of each method. DTI-CDF was originally evaluated on all three splitting schemes underlining this point (Chu *et al*., 2019). While DTI-Voodoo provides comparable performance to the naïve predictor and DTI-CDF in terms of MacroAUC, it yields considerably better results with respect to MicroAUC_*p*_. We also find that methods that are trained using a protein split generally result in higher MicroAUC_*p*_ than methods trained using a drug–target pair split, indicating that they may generalize better to unseen protein targets whereas methods trained on a drug–target split potentially exploit hidden biases and therefore generalize less well.

**Table 2.**
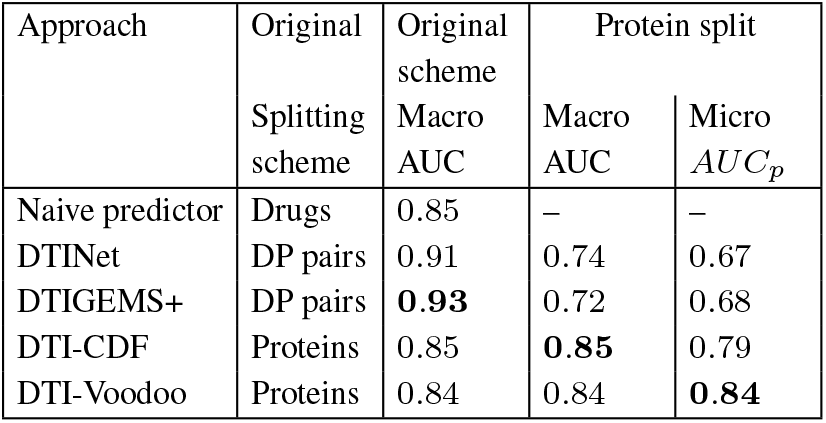
Comparison of DTI-Voodoo with state of the art drug–target interaction prediction methods on the Yamanishi dataset; we evaluated the original and the protein-based split in a cross-validation

| Approach | Original Splitting scheme | Original scheme Macro AUC | Protein split |  |
| --- | --- | --- | --- | --- |
| | | | Macro AUC | Micro $AUC_p$ |
| Naïve predictor | Drugs | 0.85 | – | – |
| DTINet | DP pairs | 0.91 | 0.74 | 0.67 |
| DTIGEMS+ | DP pairs | **0.93** | 0.72 | 0.68 |
| DTI-CDF | Proteins | 0.85 | **0.85** | 0.79 |
| DTI-Voodoo | Proteins | 0.84 | 0.84 | **0.84** |

As the difference in performance with different splitting schemes is quite large, we further evaluated additional drug–target interaction and drug–target affinity prediction methods that were trained and evaluated on other datasets. Following the results of MolTrans (Huang *et al*., 2020), we reevaluated DeepDTI (Wen *et al*., 2017), DeepDTA (Öztürk *et al*., 2018), DeepConv-DTI (Lee *et al*., 2019), and MolTrans itself on the BioSnap dataset (Zitnik *et al*., 2018) and compared it to our “naïve” predictor as well as DTI-Voodoo (see Table 3). MolTrans was evaluated over the drug– target pair and the protein split; we were able to reproduce the MolTrans results (Table 3), showing a substantial difference based on the splitting scheme. We additionally computed the *MicroAUC*_*p*_ score for all considered methods, leading to similar results as observed on the Yaminishi dataset. We test whether DTI-Voodoo performs better than the methods we compare against or whether the observed values for *MicroAUC*_*p*_ and *MacroAUC* fall within expected variant; we use a one-tailed T-test with Bonferroni correction for this test.

**Table 3.** Comparison of DTI-Voodoo with state of the art drug–target interaction prediction methods on the BioSnap dataset; we evaluated the original and the protein-based split in a cross-validation.

| Approach | Original Splitting scheme | Original scheme Macro AUC | Protein split |  |
| --- | --- | --- | --- | --- |
| | | | Macro AUC | Micro $AUC_p$ |
| Naïve predictor | DP pairs | 0.79 | – | – |
| DeepDTI | Drugs | 0.88 | 0.76 | 0.70 |
| DeepDTA | DP pairs | 0.88 | 0.77 | 0.69 |
| DeepConv-DTI | DP pairs | 0.88 | 0.76 | 0.73 |
| MolTrans | DP pairs | **0.90** | 0.77 | 0.74 |
| DTI-Voodoo | Proteins | 0.85 | **0.85** | **0.82** |

Considering Macro AUCs, DTI-Voodoo improves significantly (*p <* 10^−4^, one-sided t-test) over MolTrans (*t* = 36.6), DeepDTI (*t* = 40.5), DeepDTA (*t* = 36.6), and DeepConv-DTI (*t* = 40.5) on the BioSnap dataset, and significantly (*p <* 10^−4^, one-sided t-test) over DTINet (*t* = 20.8) and DTIGEMS+ (*t* = 18.0) on the Yamanishi dataset; DTI-CDF performs better than DTI-Voodoo (*p* = 0.1, *t* = 1.3, onesided t-test). We further perform another one-tailed t-test to compare *MicroAUC*_*p*_ performance. We find that DTI-Voodoo improves significantly (*p <* 10^−4^) over MolTrans(*t* = 21.2), DeepDTI (*t* = 31.0), DeepDTA (*t* = 33.4), and DeepConv-DTI (*t* = 23.7) on the BioSnap dataset, and over DTINet (*t* = 29.6), DTIGEMS+ (*t* = 28.0), and DTI-CDF (*t* = 9.5) on the Yamanishi dataset.

In all our experiments, DTI-Voodoo improves over all other methods with respect to *MicroAUC*_*p*_, demonstrating that DTI-Voodoo can identify drugs that target a specific protein more accurately than other methods. Some methods achieve a higher MacroAUC than DTI-Voodoo, in particular when evaluated using a drug–target pair splitting scheme; our results with the “naïve” prediction method show that it may be possible that models trained on a drug–target split utilize certain biases in the dataset without necessarily producing novel biological insights.

## 5 Discussion

### 5.1 “Bottom-up” and “top-down” prediction of drug-target interactions

There are many computational methods to predict drug–target interactions. They can broadly be grouped in two types; the first, which we refer to as “bottom-up” approaches, start from molecular information about a drug and protein and predict an interaction based on their molecular properties; the second, which we refer to as “top-down” approaches, start from observable characteristics of an organism and infer drug–target interactions as the putative molecular mechanisms that explain these observations.

Another view on these two approaches is as direct and indirect ways to predict drug–target interactions. On one hand, molecular information can be used to directly determine whether two molecules (such as a drug and protein) have the ability to interact, whereas information about phenotypic consequences of a drug (drug effects) or disruption of a protein function can be used to indirectly suggest candidate drug–target interactions. Molecular features will be specific to a drug–target pair and we would not expect this information to propagate through a protein–protein interaction network; the main information about drug–target interactions that could be obtained from interactions between proteins is information about binding sites between proteins that may also be used by a drug molecule (i.e., information that protein *P*_1_ binds to protein *P*_2_ reveals information about the molecular structures of both *P*_1_ and *P*_2_). On the other hand, phenotypic consequences of changes in protein function, or drug effects, are often a result of aberrant pathway or network activity and involve more than one protein; consequently, we expect these features to benefit more from including information about protein–protein interactions. Moreover, as the protein–protein interaction relation is not transitive (protein *P*_1_ interacting with *P*_2_, and *P*_2_ relating with *P*_3_ does not imply *P*_1_ interacting with *P*_3_), we mainly transfer information to the direct neighborhood of each protein within the PPI graph. Our results (Table 1) confirm the first hypothesis and demonstrate that molecular features do not benefit from including the interaction network whereas the indirect, top-down features benefit from the propagation over the interaction network. While the graph we chose is based on interactions between proteins and our results hold true for such a graph, other types of entities and interactions can also be chosen, in particular similarity networks (Gottlieb *et al*., 2011); in such networks, information between nodes may be transmitted differently than in DTI-Voodoo.

There are other types of indirect features that could be added to our model. A common feature that may be added are drug indications which are predictive of drug–target interactions (Gottlieb *et al*., 2011). However, we do not include them in our model as including drug indications would allow our model to make many trivial predictions based only on remembering which targets are often used for which indication; including network information would likely benefit predictions based on drug indications because different drugs may target the same pathway through different mechanisms.

Combining bottom-up and top-down approaches in a single model can follow different strategies. Interaction networks are used widely to determine indirect effects of molecular changes and predict drug–target interactions (Shahreza *et al*., 2017). Our work relies on graph neural networks as a way to combine qualitative information about interactions with additional features (molecular interaction, phenotypic and functional features); even if only some of these features benefit from the information the graph provides, graph neural networks will allow further extension of our model with additional features in the future.

To the best of our knowledge, DTI-Voodoo is the first DTI prediction model that propagates ontology-based features of protein interaction networks; while a similar approach of combining ontology embeddings with interaction networks has previously been used for analyzing gene expression (**?**), DTI-Voodoo extends this method to DTI prediction. Further, DTI-Voodoo is novel in that it explicitly integrates “bottom-up” and “top-down” features using a graph representation of interactions between proteins; while there are other DTI prediction methods that combine these features (Gottlieb *et al*., 2011), DTI-Voodoo exploits the ability to integrate heterogeneous features using graphs and the ability to utilize this graph in machine learning through the use of graph neural networks.

### 5.2 The challenge of evaluating drug–target interaction predictions

One major component of our experiments was to determine how the information that is available to a machine learning model during training affects the performance of the model. Similarly to previous work (Huang *et al*., 2020; Lee *et al*., 2019; Chu *et al*., 2019), we find significant differences in predictive performance across different splitting schemes.

The most common scheme for drug–target interaction prediction is the split over drug-target pairs (Wang and Kurgan, 2018) where it may happen that most drugs and targets that are including in the model’s validation and testing phase have also been included in the training phase (as part of other drug–target pairs). This scheme is prone to a number of biases. If the number of interactions for a drug or protein are imbalanced, i.e., some drugs or proteins have many more interactions than others, these will be seen more often during training and they will likely also have more interactions in the testing and validation sets; because some entities have more interactions, i.e., they are more likely to interact, any model that preferentially predicts these as interaction partners will improve its predictive performance. While this accurately captures the distribution, predicting based on biases in the number of interaction partners is not desirable when applying the model to novel entities. We have demonstrated that even the newly proposed “naïve” classifier that makes predictions only by exploiting the imbalanced number of interaction partners can achieve performance close to state of the art methods (when measuring Macro AUC). When training a machine learning model on such imbalanced data, it will eventually overfit to this imbalance. Splitting by entity (protein or drug) can reduce the impact of these spurious correlations but not reduce it entirely, because similar entities will still exhibit similar interaction patterns. In our experiments, we observed the impact of splitting training and evaluation sets by protein as a decrease in overall performance (Macro AUC), providing some evidence that models trained using this splitting scheme are less sensitive to overfitting to this type of bias.

The way in which training and evaluation data is generated is related to how the model is evaluated. An evaluation based on Macro AUC evaluated the application scenario where a set of drugs and proteins are given, and out of all possible pairs, the more likely interactions need to be identified. However, this does not correspond to most scenarios for drug repurposing where a drug that targets a *specific* protein (e.g., a protein involved in a disease) needs to be identified. We introduce an evaluation measure based on micro-averages per protein (Micro AUC_*p*_) to evaluate this scenario, and we often find substantial differences in predictive performance when evaluating with Macro AUC and Micro AUC_*p*_; generally, models that are trained using a split over drug–target pairs perform worse in Micro AUC_*p*_ than models that use a protein-based split, further providing evidence that a drug–target split results in overfitting to dataset biases.

Finally, a potential source of differences in model performance is how negatives are identified and treated during evaluation (and training). There are few large sets of validated negative drug–target interactions; consequently, many models (including DTI-Voodoo) use all unknown interactions as negatives. As there are many more negative than positive interactions, negatives are then sub-sampled during training resulting in a training set that is balanced between positives and negatives (or a certain picture(0,0)(−35,0)(1,0)30 (0,35)(0,-1)30 picture picture(0,0)(35,0)(−1,0)30 (0,35)(0,-1)30 picture ratio is preserved). While this is a reasonable strategy to deal with imbalanced data, it may be problematic when the same sub-sampling is applied on the model’s evaluation set because it over-simplifies the evaluation process. The performance differences is not usually visible when using ROC curves but results in unrealistically high precision and therefore high area under a precision-recall curve.

Several of the biases we identify in evaluating DTI prediction methods have been observed previously. The performance difference based on how training and evaluation data is split (by interaction pair, by drug, or by protein) has been demonstrated before using the MacroAUC measure (Huang *et al*., 2020; Pahikkala *et al*., 2014; van Laarhoven and Marchiori, 2014); we further extend on these results by introducing performance measures based on micro averages (Micro AUC_*p*_ and Micro AUC_*d*_) to further illustrate how prediction performance changes when evaluation data is imbalanced. We have further extended on prior results by introducing a “naïve” classifier that explicitly exploits one data bias to make predictions, illustrating that this bias has a significant impact on DTI methods. The bias we identify with the “naïve” classifier is similar to a previous bias found in gene networks when using methods that rely on the guilt-by-association principle (Gillis and Pavlidis, 2012) but which has, to our knowledge, not been demonstrated in the context of DTI prediction.

In summary, drug–target interaction prediction is not a single computational problem in bioinformatics but a set of related problems. Let *P* be a set of proteins *P* and *D* a set of drugs; one task can be to identify arbitrary pairs (*p, d*) with *p ∈ P* and *d ∈ D* that interact, another to identify a set of interacting drugs for each *p ∈ P*, and yet another to identify a set of proteins for each drug *d ∈ D*. The first task may be useful when no particular drug or protein is considered; the second task when searching for a drug that targets a specific (disease-associated) protein; and the third when aiming to find new applications for a given drug. The first task would best be evaluated using a Macro AUC, the second and third using a Micro AUC_*p*_ and Micro AUC_*d*_.

### 5.3 Pharmacological novelty

As the target-based predictive power of DTI-Voodoo improves significantly over other methods, we utilized our model to predict novel drug classes for protein families. We therefore collected the 2nd level ATC (Anatomical Therapeutic Chemical) groups for each drug and all InterPro (**?**) top-level families for each protein. Utilizing the STITCH interactions, we followed a protein split within each InterPro family by predicting over all available drugs. We eventually normalized the number of novel interactions per group by the amount of drugs within the respective ATC group. A heatmap showing the results of this analysis can be found in Supplementary Figure 5. DTI-Voodoo predicts novel candidate drug–target interactions for a broad range of ATC categories as well as protein families.

For example, ATC group A07 (Antidiarrheals) has relatively few approved drugs in total (Supplementary Figure 6), but DTI-Voodoo predicts several candidate targets from proteins with a PHD-type zinc finger do-main (`IPR001965`). For example, the drug mesalazine, used to treat inflammatory bowel disease but with an apoptosis-inducing and chemo-preventative effect in colon cancer (**??**); DTI-Voodoo predicts mesalazine to interact with five proteins with PHD-type zinc finger domain: BRPF1, TRIM33, BAZ1A, RSF1, and DPF2. Overexpression of RSF1 is associated with poor prognosis in colorectal cancer, and knock-down of RSF1 leads to decrease of cell proliferation (**?**), indicating that, in addition to its antiinflammatory effects, mesalazine may act through inhibition of RSF1 in its chemopreventative effects on colon cancer. We make all predictions including the ATC class of the drug and the Interpro family of the predicted target available on our website to allow further exploration of DTI-Voodoo’s prediction results.

## 6 Conclusions

We developed DTI-Voodoo as a machine learning model that combines molecular features and functional information with an interaction network using graph neural networks to predict drugs that may target specific proteins. In this task, DTI-Voodoo improves over several state of the art methods. We demonstrated that functional and phenotypic information localizes on the interaction network whereas molecular information does not. Moreover, we showed that drug–target interaction prediction datasets have some inherent biases that affect the performance of models. This led us to conclude that drug–target interaction prediction is not a single computational problem but a set of multiple problems. Experimental evaluation of drug–target interaction prediction methods must be carefully designed to reflect the problem the model aims to solve, and the interpretation of performance results should be aligned with the specific problem.

## Acknowledgements

We acknowledge the use of computational resources from the KAUST Supercomputing Core Laboratory.

## References

Ashburner, M., Ball, C. A., Blake, J. A., Botstein, D., Butler, H., Cherry, J. M., Davis, A. P., Dolinski, K., Dwight, S. S., Eppig, J. T., Harris, M. A., Hill, D. P., Issel-Tarver, L., Kasarskis, A., Lewis, S., Matese, J. C., Richardson, J. E., Ringwald, M., Rubin, G. M., and Sherlock, G. (2000). Gene ontology: tool for the unification of biology. Nature Genetics, 25(1), 25–29.

Bianchi, F. M., Grattarola, D., Livi, L., and Alippi, C. (2019). Graph neural networks with convolutional ARMA filters. CoRR, abs/1901.01343.

Campillos, M., Kuhn, M., Gavin, A.-C., Jensen, L. J., and Bork, P. (2008). Drug target identification using side-effect similarity. Science, 321(5886), 263–266.

Carbon, S., Douglass, E., Good, B. M., Unni, D. R., Harris, N. L., Mungall, C. J., Basu, S., Chisholm, R. L., Dodson, R. J., Hartline, E., Fey, P., Thomas, P. D., Albou, L.-P., Ebert, D., Kesling, M. J., Mi, H., Muruganujan, A., Huang, X., Mushayahama, T., LaBonte, S. A., Siegele, D. A., Antonazzo, G., Attrill, H., Brown, N. H., Garapati, P., Marygold, S. J., Trovisco, V., dos Santos, G., Falls, K., Tabone, C., Zhou, P., Goodman, J. L., Strelets, V. B., Thurmond, J., Garmiri, P., Ishtiaq, R., Rodríguez-López, M., Acencio, M. L., Kuiper, M., Lægreid, A., Logie, C., Lovering, R. C., Kramarz, B., Saverimuttu, S. C. C., Pinheiro, S. M., Gunn, H., Su, R., Thurlow, K. E., Chibucos, M., Giglio, M., Nadendla, S., Munro, J., Jackson, R., Duesbury, M. J., Del-Toro, N., Meldal, B. H. M., Paneerselvam, K., Perfetto, L., Porras, P., Orchard, S., Shrivastava, A., Chang, H.-Y., Finn, R. D., Mitchell, A. L., Rawlings, N. D., Richardson, L., Sangrador-Vegas, A., Blake, J. A., Christie, K. R., Dolan, M. E., Drabkin, H. J., Hill, D. P., Ni, L., Sitnikov, D. M., Harris, M. A., Oliver, S. G., Rutherford, K., Wood, V., Hayles, J., Bähler, J., Bolton, E. R., Pons, J. L. D., Dwinell, M. R., Hayman, G. T., Kaldunski, M. L., Kwitek, A. E., Laulederkind, S. J. F., Plasterer, C., Tutaj, M. A., Vedi, M., Wang, S.-J., D’Eustachio, P., Matthews, L., Balhoff, J. P., Aleksander, S. A., Alexander, M. J., Cherry, J. M., Engel, S. R., Gondwe, F., Karra, K., Miyasato, S. R., Nash, R. S., Simison, M., Skrzypek, M. S., Weng, S., Wong, E. D., Feuermann, M., Gaudet, P., Morgat, A., Bakker, E., Berardini, T. Z., Reiser, L., Subramaniam, S., Huala, E., Arighi, C. N., Auchincloss, A., Axelsen, K., Argoud-Puy, G., Bateman, A., Blatter, M.-C., Boutet, E., Bowler, E., Breuza, L., Bridge, A., Britto, R., Bye-A-Jee, H., Casas, C. C., Coudert, E., Denny, P., Estreicher, A., Famiglietti, M. L., Georghiou, G., Gos, A., Gruaz-Gumowski, N., Hatton-Ellis, E., Hulo, C., Ignatchenko, A., Jungo, F., Laiho, K., Mercier, P. L., Lieberherr, D., Lock, A., Lussi, Y., MacDougall, A., Magrane, M., Martin, M. J., Masson, P., Natale, D. A., Hyka-Nouspikel, N., Orchard, S., Pedruzzi, I., Pourcel, L., Poux, S., Pundir, S., Rivoire, C., Speretta, E., Sundaram, S., Tyagi, N., Warner, K., Zaru, R., Wu, C. H., Diehl, A. D., Chan, J. N., Grove, C., Lee, R. Y. N., Muller, H.-M., Raciti, D., Auken, K. V., Sternberg, P. W., Berriman, M., Paulini, M., Howe, K., Gao, S., Wright, A., Stein, L., Howe, D. G., Toro, S., Westerfield, M., Jaiswal, P., Cooper, L., and Elser, J. (2020). The gene ontology resource: enriching a GOld mine. Nucleic Acids Research, 49(D1), D325–D334.

Chen, J., Althagafi, A., and Hoehndorf, R. (2020). Predicting candidate genes from phenotypes, functions and anatomical site of expression. Bioinformatics. advance access.

Chen, X., Yan, C. C., Zhang, X., Zhang, X., Dai, F., Yin, J., and Zhang, Y. (2015). Drug–target interaction prediction: databases, web servers and computational models. Briefings in Bioinformatics, 17(4), 696–712.

Chu, Y., Kaushik, A. C., Wang, X., Wang, W., Zhang, Y., Shan, X., Salahub, D. R., Xiong, Y., and Wei, D.-Q. (2019). DTI-CDF: a cascade deep forest model towards the prediction of drug-target interactions based on hybrid features. Briefings in Bioinformatics, 22(1), 451–462.

Defferrard, M., Bresson, X., and Vandergheynst, P. (2016). Convolutional neural networks on graphs with fast localized spectral filtering. In Proceedings of the 30th International Conference on Neural Information Processing Systems, NIPS’16, page 3844–3852, Red Hook, NY, USA. Curran Associates Inc.

Ding, H., Takigawa, I., Mamitsuka, H., and Zhu, S. (2013). Similarity-based machine learning methods for predicting drug–target interactions: a brief review. Briefings in Bioinformatics, 15(5), 734–747.

Feng, Y., Wang, Q., and Wang, T. (2017). Drug target protein-protein interaction networks: A systematic perspective. BioMed Research International, 2017, 1–13.

Fey, M. and Lenssen, J. E. (2019). Fast graph representation learning with pytorch geometric. CoRR, abs/1903.02428.

Gillis, J. and Pavlidis, P. (2012). “guilt by association” is the exception rather than the rule in gene networks. PLoS Computational Biology, 8(3), e1002444.

Gottlieb, A., Stein, G. Y., Ruppin, E., and Sharan, R. (2011). PREDICT: a method for inferring novel drug indications with application to personalized medicine. Molecular Systems Biology, 7(1), 496.

Hamilton, W. L., Ying, Z., and Leskovec, J. (2017). Inductive representation learning on large graphs. In NIPS.

Hoehndorf, R., Schofield, P. N., and Gkoutos, G. V. (2011). PhenomeNET: a whole-phenome approach to disease gene discovery. Nucleic Acids Research, 39(18), e119–e119.

Honda, S., Shi, S., and Ueda, H. R. (2019). SMILES transformer: Pre-trained molecular fingerprint for low data drug discovery. CoRR, abs/1911.04738.

Huang, K., Xiao, C., Glass, L. M., and Sun, J. (2020). MolTrans: Molecular interaction transformer for drug–target interaction prediction. Bioinformatics. advanced access.

Jeni, L. A., Cohn, J. F., and De La Torre, F. (2013). Facing imbalanced data– recommendations for the use of performance metrics. In 2013 Humaine Association Conference on Affective Computing and Intelligent Interaction, pages 245–251.

Kingma, D. P. and Ba, J. (2015). Adam: A method for stochastic optimization. CoRR, abs/1412.6980.

Kipf, T. N. and Welling, M. (2016). Semi-supervised classification with graph convolutional networks. CoRR, abs/1609.02907.

Klicpera, J., Bojchevski, A., and Günnemann, S. (2018). Personalized embedding propagation: Combining neural networks on graphs with personalized pagerank. CoRR, abs/1810.05997.

Köhler, S., Carmody, L., Vasilevsky, N., Jacobsen, J. O. B., Danis, D., Gourdine, J.-P., Gargano, M., Harris, N. L., Matentzoglu, N., McMurry, J. A., Osumi-Sutherland, D., Cipriani, V., Balhoff, J. P., Conlin, T., Blau, H., Baynam, G., Palmer, R., Gratian, D., Dawkins, H., Segal, M., Jansen, A. C., Muaz, A., Chang, W. H., Bergerson, J., Laulederkind, S. J. F., Yüksel, Z., Beltran, S., Freeman, A. F., Sergouniotis, P. I., Durkin, D., Storm, A. L., Hanauer, M., Brudno, M., Bello, S. M., Sincan, M., Rageth, K., Wheeler, M. T., Oegema, R., Lourghi, H., Rocca, M. G. D., Thompson, R., Castellanos, F., Priest, J., Cunningham-Rundles, C., Hegde, A., Lovering, R. C., Hajek, C., Olry, A., Notarangelo, L., Similuk, M., Zhang, X. A., Gómez-Andrés, D., Lochmüller, H., Dollfus, H., Rosenzweig, S., Marwaha, S., Rath, A., Sullivan, K., Smith, C., Milner, J. D., Leroux, D., Boerkoel, C. F., Klion, A., Carter, M. C., Groza, T., Smedley, D., Haendel, M. A., Mungall, C., and Robinson, P. N. (2018). Expansion of the human phenotype ontology (HPO) knowledge base and resources. Nucleic Acids Research, 47(D1), D1018–D1027.

Kuhn, M., Letunic, I., Jensen, L. J., and Bork, P. (2015). The SIDER database of drugs and side effects. Nucleic Acids Research, 44(D1), D1075–D1079.

Kulmanov, M. and Hoehndorf, R. (2019). DeepGOPlus: improved protein function prediction from sequence. Bioinformatics, 36(2), 422–429.

Lee, I. and Nam, H. (2018). Identification of drug-target interaction by a random walk with restart method on an interactome network. BMC Bioinformatics, 19(S8).

Lee, I., Keum, J., and Nam, H. (2019). DeepConv-DTI: Prediction of drug-target interactions via deep learning with convolution on protein sequences. PLOS Computational Biology, 15(6), e1007129.

Li, G., Müller, M., Thabet, A., and Ghanem, B. (2019). Deepgcns: Can gcns go as deep as cnns? In The IEEE International Conference on Computer Vision (ICCV).

Li, G., Xiong, C., Thabet, A., and Ghanem, B. (2020). Deepergcn: All you need to train deeper gcns. CoRR, abs/2006.07739.

Liu, Q., Hu, Z., Jiang, R., and Zhou, M. (2020). DeepCDR: a hybrid graph convolutional network for predicting cancer drug response. Bioinformatics, 36(Supplement_2), i911–i918.

Luo, Y., Zhao, X., Zhou, J., Yang, J., Zhang, Y., Kuang, W., Peng, J., Chen, L., and Zeng, J. (2017). A network integration approach for drug-target interaction prediction and computational drug repositioning from heterogeneous information. Nature communications, 8(1), 1–13.

Mikolov, T., Sutskever, I., Chen, K., Corrado, G., and Dean, J. (2013). Distributed representations of words and phrases and their compositionality. CoRR, abs/1310.4546.

Mozzicato, P. (2009). MedDRA. Pharmaceutical Medicine, 23(2), 65–75.

Nguyen, T., Le, H., Quinn, T. P., Nguyen, T., Le, T. D., and Venkatesh, S. (2020). GraphDTA: Predicting drug–target binding affinity with graph neural networks. Bioinformatics. advance access.

Oliver, S. (2000). Guilt-by-association goes global. Nature, 403(6770), 601–602.

Overington, J. P., Al-Lazikani, B., and Hopkins, A. L. (2006). How many drug targets are there? Nature Reviews Drug Discovery, 5(12), 993–996.

Öztürk, H., Özgür, A., and Ozkirimli, E. (2018). DeepDTA: deep drug–target binding affinity prediction. Bioinformatics, 34(17), i821–i829.

Pahikkala, T., Airola, A., Pietila, S., Shakyawar, S., Szwajda, A., Tang, J., and Aittokallio, T. (2014). Toward more realistic drug-target interaction predictions. Briefings in Bioinformatics, 16(2), 325–337.

Shahreza, M. L., Ghadiri, N., Mousavi, S. R., Varshosaz, J., and Green, J. R. (2017). A review of network-based approaches to drug repositioning. Briefings in Bioinformatics, 19(5), 878–892.

Smith, C. L. and Eppig, J. T. (2009). The mammalian phenotype ontology: enabling robust annotation and comparative analysis. Wiley Interdisciplinary Reviews: Systems Biology and Medicine, 1(3), 390–399.

Szklarczyk, D., Franceschini, A., Wyder, S., Forslund, K., Heller, D., Huerta-Cepas, J., Simonovic, M., Roth, A., Santos, A., Tsafou, K. P., Kuhn, M., Peer Jensen, L.J., and von Mering, C. (2014). STRING v10: protein–protein interaction networks, integrated over the tree of life. Nucleic Acids Research, 43(D1), D447–D452.

Szklarczyk, D., Santos, A., von Mering, C., Jensen, L. J., Bork, P., and Kuhn, M. (2015). STITCH 5: augmenting protein–chemical interaction networks with tissue and affinity data. Nucleic Acids Research, 44(D1), D380–D384.

Thafar, M. A., Olayan, R. S., Ashoor, H., Albaradei, S., Bajic, V. B., Gao, X., Gojobori, T., and Essack, M. (2020). DTiGEMS: drug-target interaction prediction using graph embedding, graph mining, and similarity-based techniques. Journal of Cheminformatics, 12(1).

van Laarhoven, T. and Marchiori, E. (2014). Biases of drug–target interaction network data. In Pattern Recognition in Bioinformatics, pages 23–33. Springer International Publishing.

Veličković, P., Cucurull, G., Casanova, A., Romero, A., Lio, P., and Bengio, Y. (2017). Graph attention networks. CoRR, abs/1710.10903v3.

Wan, F., Hong, L., Xiao, A., Jiang, T., and Zeng, J. (2019). NeoDTI: neural integration of neighbor information from a heterogeneous network for discovering new drug–target interactions. Bioinformatics, 35(1), 104–111.

Wang, C. and Kurgan, L. (2018). Review and comparative assessment of similarity-based methods for prediction of drug–protein interactions in the druggable human proteome. Briefings in Bioinformatics, 20(6), 2066–2087.

Wen, M., Zhang, Z., Niu, S., Sha, H., Yang, R., Yun, Y., and Lu, H. (2017). Deeplearning-based drug–target interaction prediction. Journal of Proteome Research, 16(4), 1401–1409.

Wishart, D. S., Knox, C., Guo, A. C., Cheng, D., Shrivastava, S., Tzur, D., Gautam, B., and Hassanali, M. (2007). DrugBank: a knowledgebase for drugs, drug actions and drug targets. Nucleic Acids Research, 36(Suppl_1), D901–D906.

Wishart, D. S., Feunang, Y. D., Guo, A. C., Lo, E. J., Marcu, A., Grant, J. R., Sajed, T., Johnson, D., Li, C., Sayeeda, Z., Assempour, N., Iynkkaran, I., Liu, Y., Maciejewski, A., Gale, N., Wilson, A., Chin, L., Cummings, R., Le, D., Pon, A., Knox, C., and Wilson, M. (2017). DrugBank 5.0: a major update to the DrugBank database for 2018. Nucleic Acids Research, 46(D1), D1074–D1082.

Yamanishi, Y., Araki, M., Gutteridge, A., Honda, W., and Kanehisa, M. (2008). Prediction of drug-target interaction networks from the integration of chemical and genomic spaces. Bioinformatics, 24(13), i232–i240.

Zitnik, M. and Leskovec, J. (2017). Predicting multicellular function through multilayer tissue networks. CoRR, abs/1707.04638.

Zitnik, M., Sosic, R., and Leskovec, J. (2018). Biosnap datasets: Stanford biomedical network dataset collection. http://snap.stanford.edu/biodata, 5(1).

