## Supplemental tables and figures for "DTI-Voodoo: machine learning over interaction networks and ontology-based background knowledge predicts drug–target interactions"

### DTI-Voodoo: Supplement data

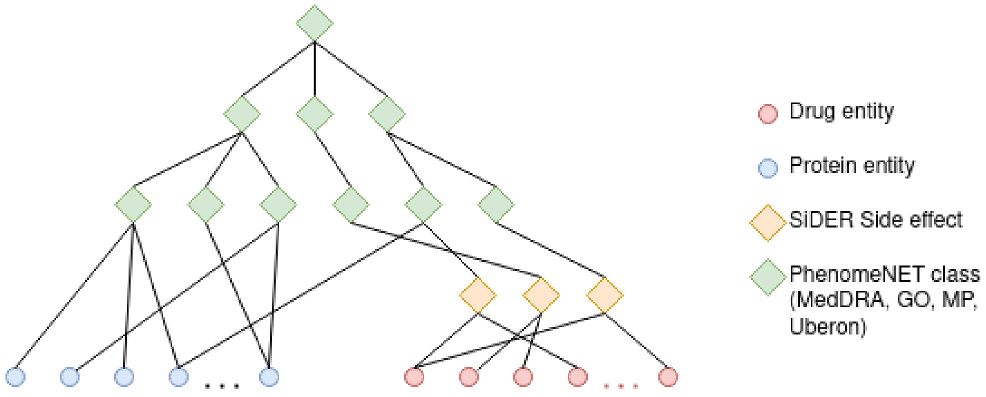


Figure 1: Drugs and proteins with annotations to SiDER and PhenomeNET

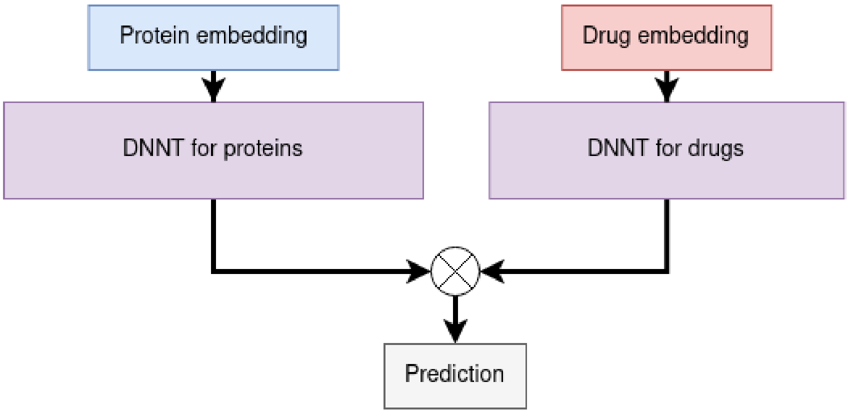


Figure 2: Half-twin network applied to molecular and DL2vec features, utilizing deep learnable feature transformations (LFT). The similarity function $\otimes$ yields the similarity between both transformed embeddings e.g. by computing the cosine similarity.

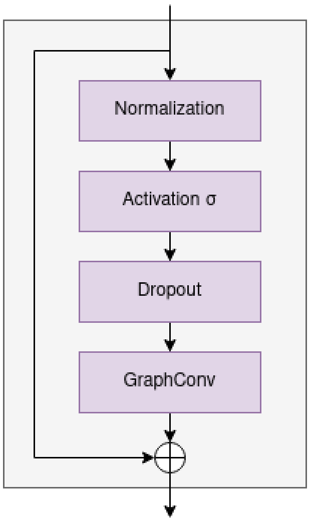


Figure 3: Residual architecture built by *Li et al. (2019, 2020b)* enabling deeper graph convolutional models

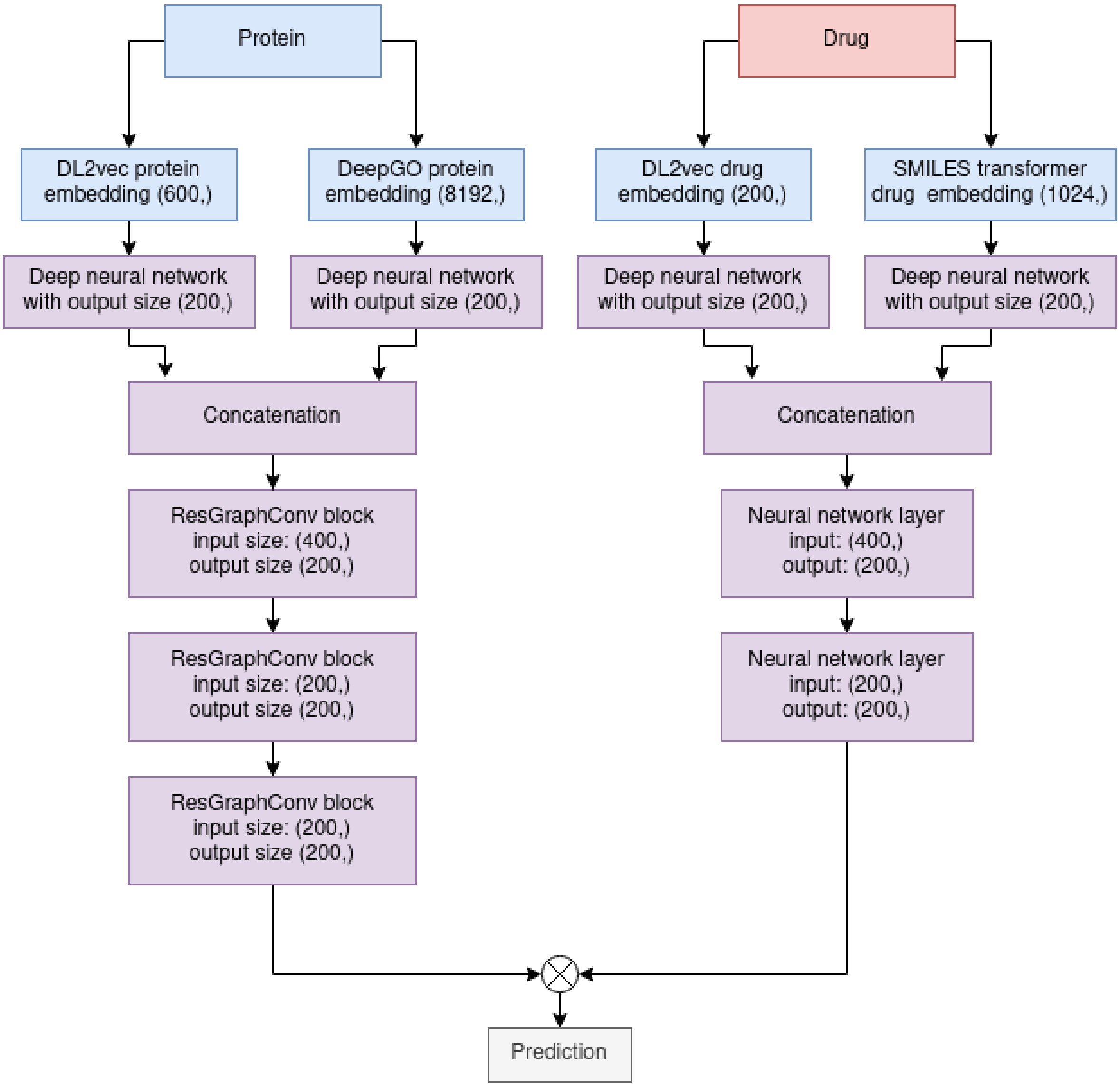


Figure 4: Residual architecture built by *Li et al. (2019, 2020b)* enabling deeper graph convolutional models

| | (a) STITCH results | | | | | |
|---|---|---|---|---|---|---|
| DTI-Voodoo results | PPI graph | | | | | |
| | without | | | with | | |
| | Macro AUC | Micro $AUC'_p$ | Micro $AUC_p$ | Macro AUC | Micro $AUC'_p$ | Micro $AUC_p$ |
| MolPred | 0.69 | 0.66 | 0.65 | 0.69 | 0.68 | 0.67 |
| OntoPred | 0.88 | 0.89 | 0.87 | 0.92 | 0.94 | 0.93 |
| DTI-Voodoo | 0.89 | 0.91 | 0.90 | 0.93 | 0.94 | 0.94 |

Table 1: Extended results table for MolPred, OntoPred and DTI-Voodoo over STITCH with added $MicroAUC'_p$ (without imputation) for comparison of both metrics.

| Approach | Graph convolution method (MacroAUC) | | | | | |
|---|---|---|---|---|---|---|
| | without | without/ empty RGC | GCN-Conv | GEN-Conv | RGC + GCN-Conv | RGC + GEN-Conv |
| MolPred | 0.69 | 0.67 | 0.67 | 0.68 | 0.65 | 0.69 |
| OntoPred | 0.88 | 0.87 | 0.88 | 0.90 | 0.86 | 0.92 |
| DTI-Voodoo | 0.89 | 0.89 | 0.89 | 0.91 | 0.86 | **0.93** |

Table 2: Results for various neural graph convolutional methods enhancing MolPred, OntoPred and DTI-Voodoo over the STITCH dataset. We evaluated all methods over different stacking heights without and with residual graph convolution blocks (RGC, ResGraphConv, see Supplementary figure 3). "Empty RGC" denotes a RGC block with a plain neural layer substituting the GraphConv layer.

| | Interaction type | | | |
|---|---|---|---|---|
| | Activation | | Inhibition | |
| #Drugs | 723 | | 954 | |
| #Proteins | 4562 | | 6222 | |
| #Links | 14174 | | 16364 | |
| Metric | Macro AUC | Micro $AUC_p$ | Macro AUC | Micro $AUC_p$ |
| DTI-Voodoo | 0.95 | 0.95 | 0.93 | 0.94 |

Table 3: Extended results table for MolPred, OntoPred and DTI-Voodoo over STITCH with added $MicroAUC'_p$ (without imputation) for comparison of both metrics.

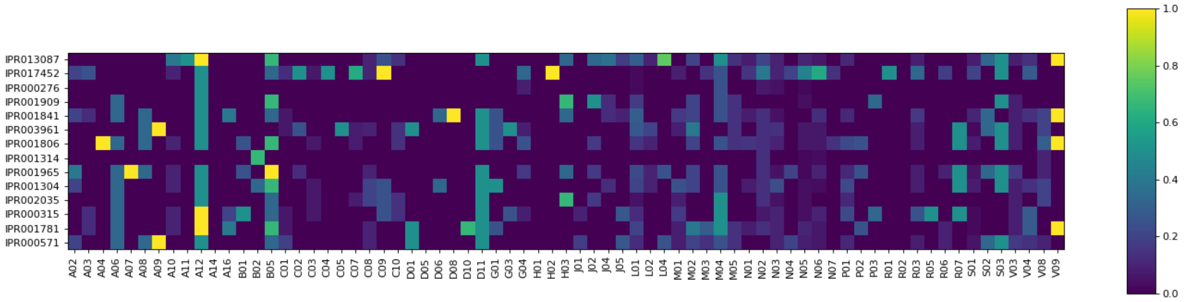


Figure 5: Heatmap showing the results of the ATC drug class and InterPro family prediction analysis. The figure shows the proportion of false negatives compared to the total amount of drugs in the corresponding level 2 ATC class. While few ATC classes show a high share of novel interactors, this is mainly due to the total amount of drugs in the ATC class. For more information note the correspondence with Supplementary Figure 6.

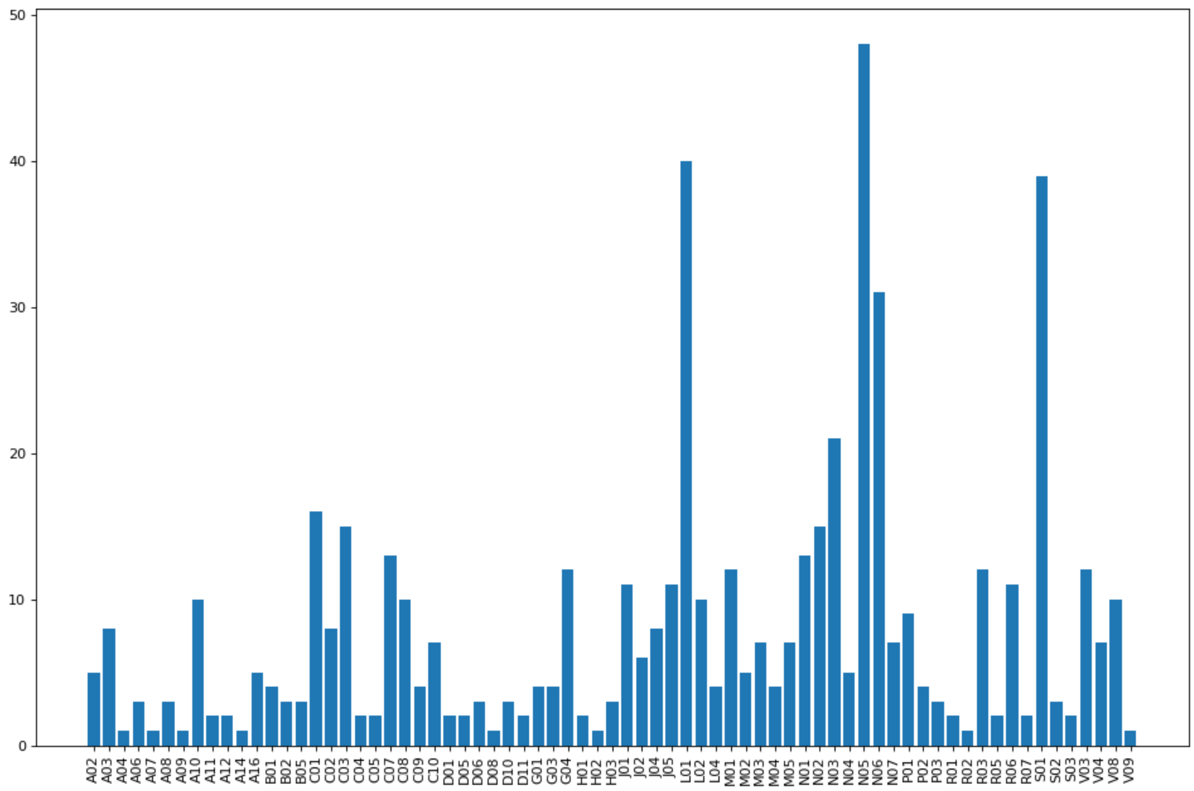


Figure 6: Amount of drugs used in the ATC/InterPro analysis for each level 2 ATC class. Notably, some of the classes have only few or single drug members.
